# A network of transcriptional repressors mediates auxin response specificity

**DOI:** 10.1101/448860

**Authors:** Jekaterina Truskina, Jingyi Han, Carlos S. Galvan-Ampudia, Stéphanie Lainé, Géraldine Brunoud, Silvana Porco, Anne-Maarit Bågman, Margot E. Smit, Malcolm Bennett, Francois Roudier, Siobhan M. Brady, Anthony Bishopp, Teva Vernoux

## Abstract

The regulation of signalling capacity plays a pivotal role in setting developmental patterns in both plants and animals (1). The hormone auxin is a key signal for plant growth and development that acts through the AUXIN RESPONSE FACTOR (ARF) transcription factors (2). A subset of these ARFs comprises transcriptional activators of target genes in response to auxin, and are essential for regulating auxin signalling throughout the plant lifecycle (3). While ARF activators show tissue-specific expression patterns, it is unknown how their expression patterns are established. Chromatin modifications and accessibility studies revealed the chromatin of loci encoding ARF activators is constitutively open for transcription. Using a high-throughput yeast one-hybrid (Y1H) approach, we discovered a network of transcriptional regulators of *ARF* activator genes from *Arabidopsis thaliana*. Expression analyses demonstrated that the majority of these regulators act as repressors of ARF transcription *in planta*. Our observations support a scenario where the default configuration of open chromatin enables a network of transcriptional repressors to shape the expression pattern of ARF activators and provide specificity in auxin signalling output throughout development.

## MAIN TEXT

*Arabidopsis thaliana* encodes 23 ARFs that act as auxin signalling effectors and trigger cellular reprogramming (2, 3), but only ARF5, 6, 7, 8 and 19 function as activators of transcription (4, 5). These five ARFs constitute the conserved class A of ARFs in Arabidopsis (abbreviated ARF^ClassA^) (6). Loss of function mutations in any of these loci can have drastic effects on embryonic and post-embryonic development. ARF^ClassA^ members are essential for the establishment and activity of root and the shoot apical meristems (RAM and SAM) (7); tissues that contain the stem cell niches driving post-embryonic plant development. They have been involved in embryonic root and shoot formation (ARF5), lateral branching (ARF5, 6, 7, 8 and 19), as well as tropisms (ARF7 and 19) (7-14). ARF^ClassA^ members have been shown to regulate different target genes (10, 11, 13). The tissue-specific variation of ARF^ClassA^ expression observed in both the RAM and SAM is also thought to drive specificity in downstream signalling and underpin the diversity of auxin responses (Fig. 1 a-j) (15, 16). Divergent patterns of *ARF*^*ClassA*^ expression could be due to tissue-specific differences in chromatin accessibility of their loci. They could also arise from regulatory networks comprising distinct sets of regulatory transcription factors (TFs); and in both prokaryotes and eukaryotes such networks typically incorporate a combination of transcriptional activators and repressors. All five ARF^ClassA^ are encoded by loci with a similar structure; coding sequences are interrupted by 11-14 introns, and, in the case of ARF5, 7 and 19, the first intron is 2-3 times bigger than other introns (Fig. S1a). To identify the regulatory sequences establishing ARF patterns, we tested the role of both upstream sequences (17) and the first intron (18) in determining *ARF*^*ClassA*^expression by comparing patterns from transcriptional reporter lines using either sequences 3-5 kb 5’ of the ATG and 3’ up to the end of the first intron or the 5’ sequences alone (Fig. 1k). A difference between the two reports was seen only for *ARF7*: the transcriptional reporters that included the first intron showed strong expression in the RAM (Fig. 1c), but this pattern was dramatically reduced when the sequence encompassing the first intron is missing (Fig. 1l, Fig. S1). We therefore conclude that the 3’ sequence including the intron contains regulatory information required for *ARF7* expression. In the root, the expression patterns of *ARF*^*ClassA*^members differed from previously published patterns obtained with shorter 2 kb (5’ of ATG) promoters (Fig. 1 a-e, (15)). In the shoot, we also found *ARF5* and *6* reporters recapitulate the patterns observed with RNA *in situ* hybridization more accurately than those with 2 kb promoters (Fig. 1 f-j, Fig. S2 l-p, (16)).

**Fig. 1.**
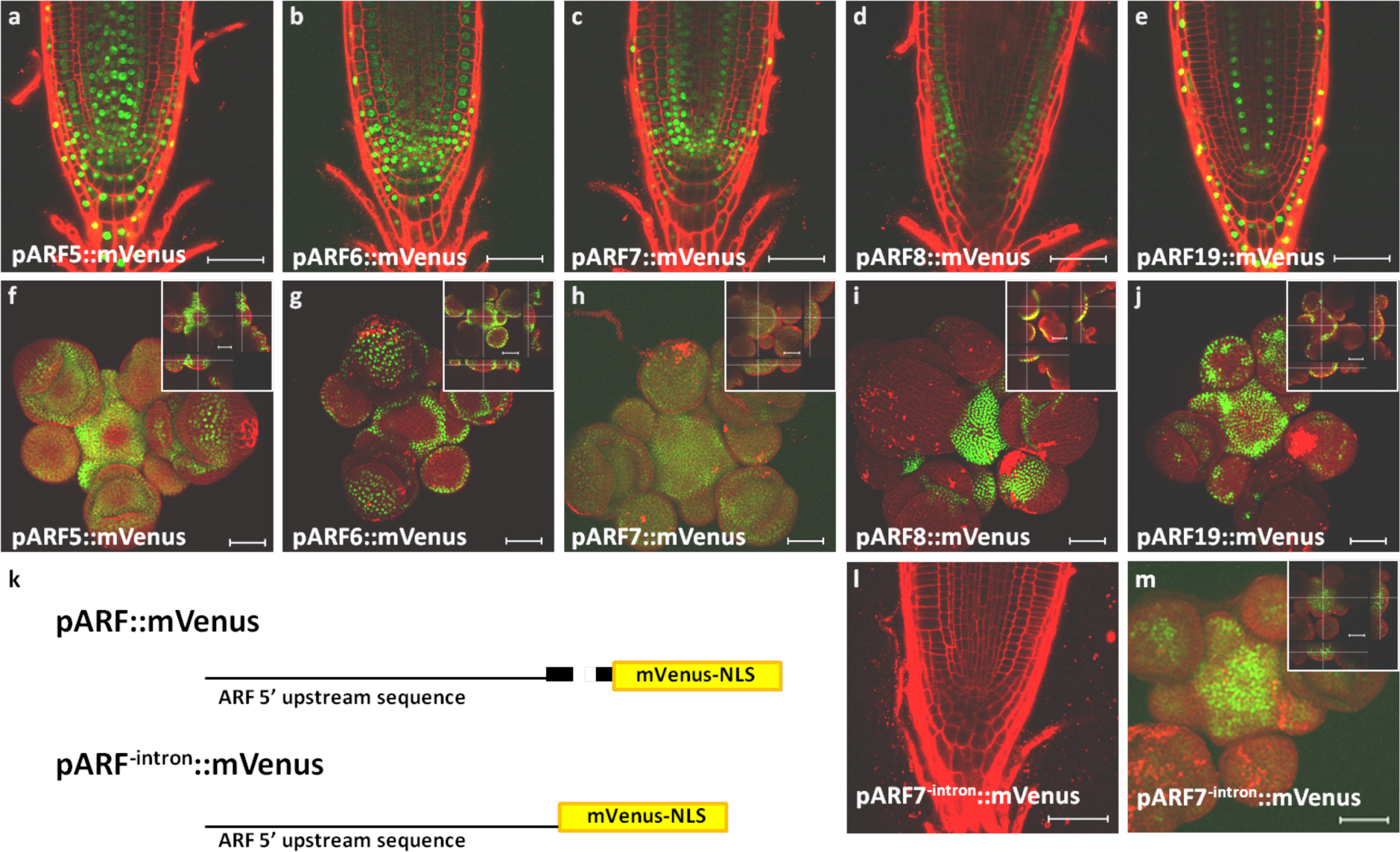
Arabidopsis Class A ARFs show tissue specific expression patterns in both the RAM and the SAM. Confocal microscopy images showing expression of *ARF5* (a, f), *ARF6* (b, g), *ARF7* (c, h), *ARF8* (d, i) and *ARF19* (e, j) in the RAM and the SAM reported using long promoters containing both 5’ and 3’ of the ATG including the first intron (*pARF::mVenus*). (k) Schematic representations of the different reporter constructs, *pARF::mVenus* and *pARF ^-intron^::mVenus*. Exons are shown in black. Expression of ARF7 in the RAM (l) and the SAM (m) reported using promoter lacking downstream sequences (k bottom). Scale bar 50 µm.

To identify the factors which determine these specific ARF expression patterns, we first analyzed the chromatin status of each *ARF*^*ClassA*^locus by assaying for H3K27me3 and H3K4me3 chromatin modifications, as these are implicated in repressing and promoting gene expression, respectively (19). Meta-analysis of published datasets covering a range of tissues and developmental stages (including the RAM and SAM) revealed H3K27me3 is largely absent from *ARF5, 6, 7, 8* and *19* loci, whereas H3K4me3 is detected at these loci (Fig. 2 a-b, Fig. S2, Supplementary Table 1). Accessible regulatory regions also characterize *ARF*^*ClassA*^ loci in the majority of tissues (Fig. 2 a-b, Fig. S3, Supplementary Table 1). These properties suggest that the chromatin configuration of *ARF*^*ClassA*^ loci allows them to be actively transcribed throughout development.

**Fig. 2.**
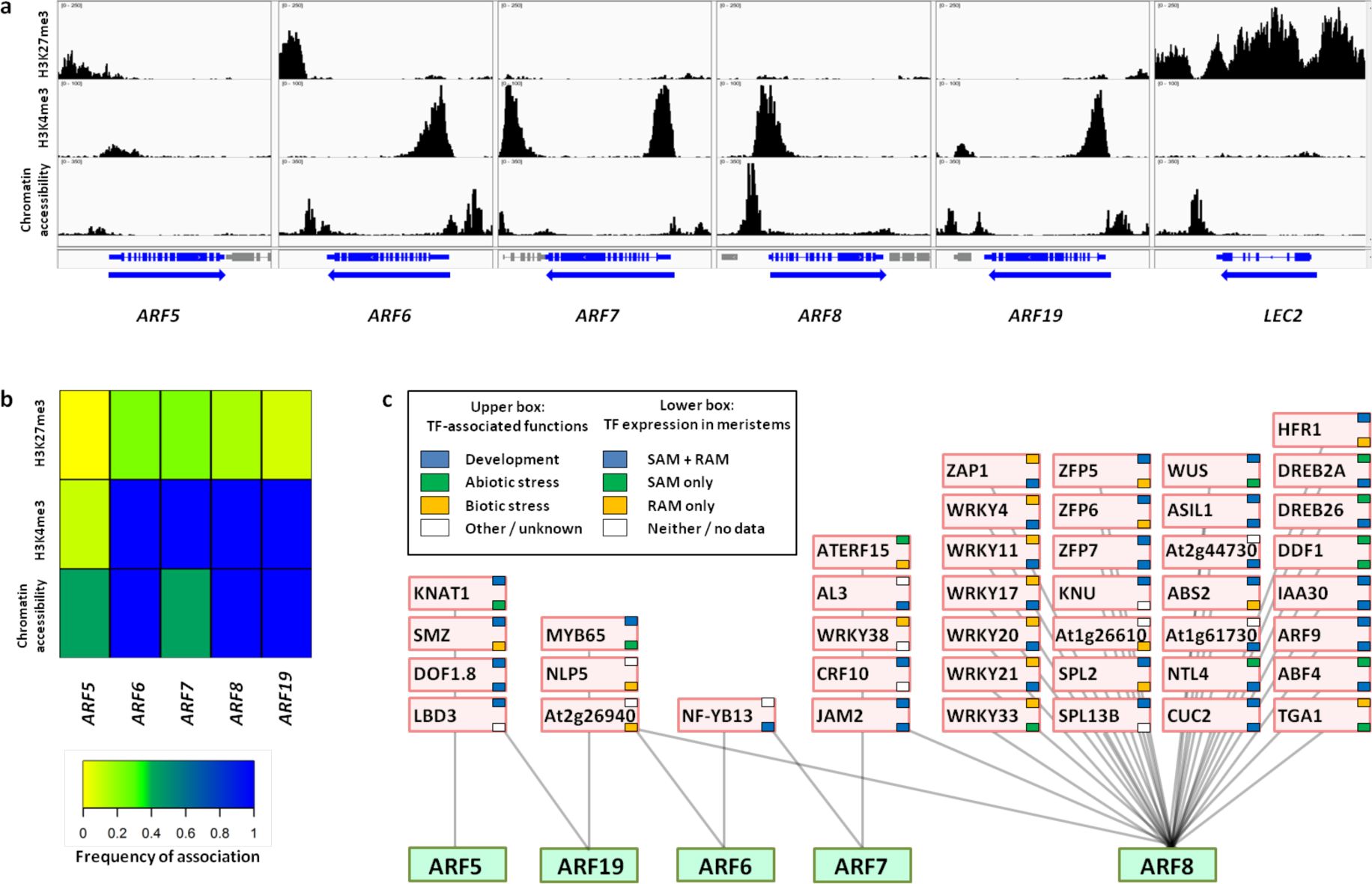
Epigenetic and transcription factor-mediated regulation of class A ARFs. (a) Chromatin landscape of ARF^ClassA^ and LEC2 loci showing distribution of the repressive H3K27me3 marker (top row) (20), the active H3K4me3 marker (middle row) (20) and the FANS-ATAC chromatin accessibility (bottom row) (21). Gene models are shown at the bottom with arrows indicating direction of transcription. LEC2 gene is included as a negative control, as it is marked by the repressive chromatin marker H3K27me3 (22). (b) Frequency of association of the repressive chromatin marker H3K27me3, active chromatin marker H3K4me3 and chromatin accessibility with the ARF^ClassA^ loci across all available datasets (see Supplementary Table 1). (c) Yeast one-hybrid promoter-transcription factor interaction network for ARF^ClassA^. Green boxes correspond to the ARF^ClassA^; pink boxes are transcription factors binding to the ARFs. TF-associated functions and expression analysis are indicated in the upper and lower small boxes and color-coded as indicated in the key.

To identify individual TFs that could regulate spatiotemporal transcription of *ARF*^*ClassA*^, we used a semi-automated, enhanced Y1H (eY1H) assay (23) with baits covering the long promoter sequences (including the first intron) identical to those from the transcriptional reporter lines described above. To do so, an existing prey collection enriched in root-stele expressed TFs was expanded to include TFs acting in the SAM and hormone signalling pathways. This extended the library’s capacity to identify putative regulators in both the root and shoot tissues, plus crosstalk between hormonal pathways. In total, this eY1H screen identified 42 novel putative transcriptional regulators of *ARF*^*ClassA*^. Mapping this gene regulatory network revealed that individual *ARF*^*ClassA*^ are regulated by distinct sets of largely non-overlapping TFs, with only 4 binding to multiple *ARF*^*ClassA*^ sequences (Fig. 2c). This network is overrepresented by members of the WRKY, ZFP, AP2/ERF and SPL TF families and contain proteins that mediate either root- or shoot-specific responses; 50% of the identified TFs are expressed in both shoots and roots while 24% and 14% are expressed specifically in roots or shoots respectively (Fig. S4 a and b). The majority of the TFs from the network are involved in development, but many putative regulators of ARF8 are associated with biotic and abiotic stress (Fig. S4c, Supplementary Table 2). This suggests ARF8 may act as an environmental hub to mediate auxin responsiveness.

To validate the *ARF*^*ClassA*^regulatory network, we first used DNA affinity purification sequencing (DAP-Seq) datasets to analyse direct binding *in vitro* of the putative regulator TFs on ARF regulatory regions (24). DAP-Seq data are available for 17 TFs and 9 (53%) of them show at least one specific peak on *ARF*^*ClassA*^promoters (Fig. S5 a and b). These analyses demonstrate binding of a significant subset of the regulatory TFs to *ARF*^*ClassA*^promoters. A literature search identified binding sites for a further 9 TFs absent from the DAP-seq dataset, supporting a model of direct regulation of *ARF^Clas^sA* by these TFs (Fig. S5c, Supplementary Table 3). We next tested the activity of each TF in the ARF^ClassA^ regulatory network systematically by employing transient expression assays in *Arabidopsis*protoplasts (Fig. 3a and b, Fig. S6). Each interaction was analysed in at least 4 independent experiments using either the TF alone or a fusion of the TF to a VP16 transactivation domain (Supplementary Table. 4). Transient expression of 27 out of 42 (64%) TFs resulted in a significant change in the expression of the *ARF*^*ClassA*^target, corresponding to a decrease in *ARF*^*ClassA*^mRNA level in 23 out of 27 of the cases (85 %; 54% of total number of TFs) (Fig. 3c). Repression of *ARF*^*ClassA*^transcription was frequently observed for both TFs alone and TF-VP16 fusions, indicating a strong repressive activity (Fig. 3b). Taken together these data revealed that the functional regulatory networks controlling *ARF*^*ClassA*^ transcription are mostly regulated by repression.

**Fig. 3.**
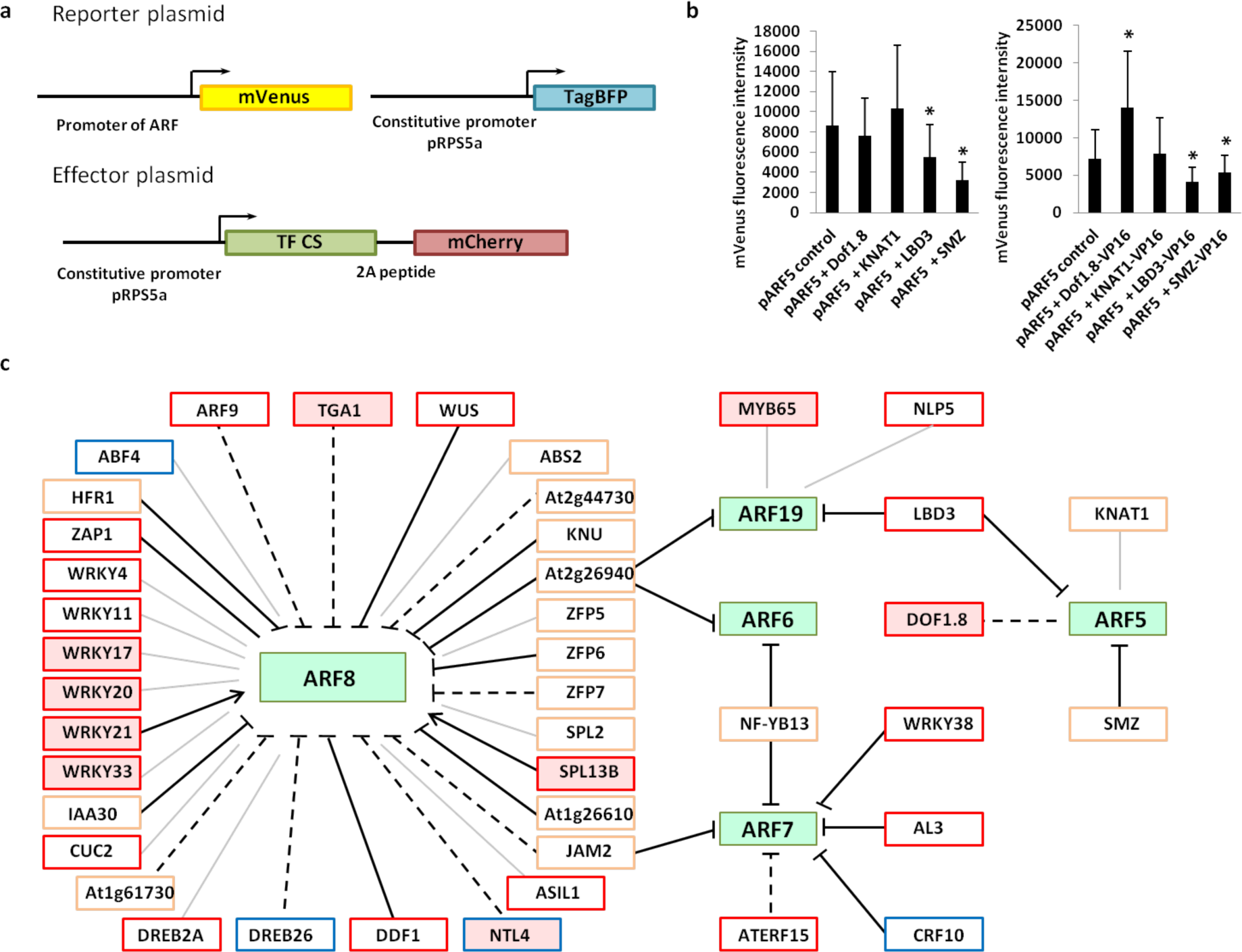
Class A ARF transcription is regulated by repressors. Interactions between ARF^ClassA^ promoters and the regulatory TFs were confirmed using transient protoplast assay. (a) Schematic of the reporter and effector plasmids co-transformed in the transient protoplast assay. (b) An example of a result from a protoplast assay using the ARF5 reporter plasmid. Two types of effector plasmids were used: one with and one without a VP16 activator domain fused to the TF coding sequence (left and right). An unpaired t-test was conducted with p < 0.1 considered as statistically significant. (c) Yeast one-hybrid promoter-transcription factor interaction network for ARF^ClassA^. Green boxes correspond to the ARF^ClassA^. TFs for which binding has been shown by DAP-seq are shown with a light red background (24) (see Supplementary Table 3). Red boarders indicate the presence of putative TF binding sites in the promoter based on published literature; blue borders indicate examples where no putative binding cites were found in the promoter. For some loci, there is insufficient data to determine binding sites, and these are shown with an orange border. Solid lines indicate interactions (arrows: activation; bars: repression) that were confirmed in the transient protoplast assay with score 4 or 3, dashed lines with score 2, and thin grey line with score 1 or 0 (see Supplementary Table 4).

To confirm these results *in planta*, we identified 25 mutants in TFs from the regulatory network representing regulators of all five ARF^ClassA^genes (Supplementary Table 5). We then measured the expression of target *ARF*^*ClassA*^genes using qRT-PCR in whole-root and shoot tissues (Fig. S7a and b, Supplementary Table 6). These detected expression changes in *ARF*^*ClassA*^targets in 11 out of 25 mutants (44%). Four showed up-regulation of their target ARFs compatible with a repressive activity. The other seven, six of which are ARF8 regulators, showed a down-regulation of their target ARF. This could be explained by a complex, non-linear regulation of ARF8 (Fig. S8) or by secondary effects of mutations. The low-sensitivity of expression analysis on whole tissues prompted us to determine at higher spatial resolution how these mutant alleles affect *ARF*^*ClassA*^expression. We focused on ARF7 and crossed the *pARF7::mVenus* transcriptional reporter into a number of mutants to examine the spatial domain of *ARF7* expression in more detail. For two of the regulator mutant backgrounds in which we had not observed changes in *ARF7* mRNA levels using qRT-PCR (*crf10* and *wrky38*), we observed a significant expansion of *pARF7::mVenus* expression pattern in the RAM (Fig. S7c and d). For *nf-yb13* we also observed enhanced expression in the RAM similar to qRT-PCR assays (Fig. 4a). This confirms *in planta* that these three TFs are repressors of *ARF*^*ClassA*^and provides an illustration of how repressors can shape *ARF* expression pattern.

**Fig. 4.**
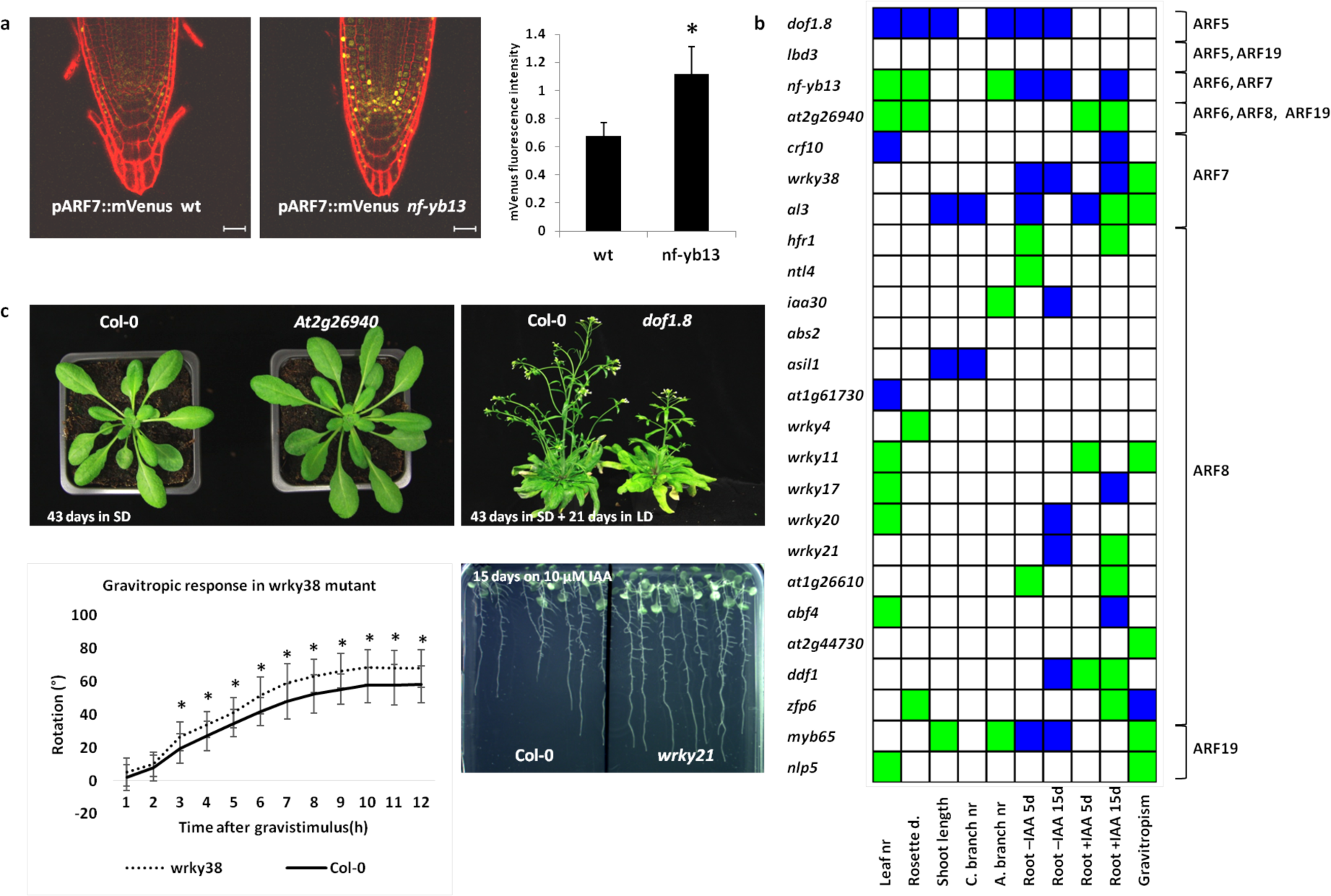
Transcriptional regulation of class A ARFs regulates a variety of developmental processes. (a) Expression of ARF7 in the RAM of *nfyb13* mutant (n= 23 wild-type and 25 mutant plants). An unpaired t-test was used to calculate significance with p ≤ 0.0001. Scale bar 25 µm. (b) Phenotypic analysis of the shoot and the root growth defects in TF mutants. Leaf nr = leaf number, Rosette d. = Rosette diameter, C. branch nr = cauline branch number, A. branch nr = axillary branch number, root = root length of plants grown with or without exogenous auxin (IAA) at 5 or 15 days, Gravitropism = gravitropic assay. Green boxes indicate statistically significant increases, blue boxes indicate statistically significant reductions in indicated growth parameter compared to Col-0. For the root growth measurements and the gravitropic response statistical analyses were done with one-way analysis of variance (ANOVA) test with p ≤ 0.05 considered as statistically significant. For the shoot growth measurements, statistical analyses were done with test an unpaired t-test with p ≤ 0.05 considered as statistically significant. (c) Examples of root and shoot growth phenotypes: shoot growth during vegetative stage in the *at2g26940*mutant (upper left) and during reproductive stage in the *dof1.8* mutant (upper right), gravitropic response in *wrky38* mutant (lower left) and root growth on 10 µM IAA at 15 days in *wrky21* (lower right).

To further confirm a role for this network in *ARF*^*ClassA*^tissue-specific repression *in planta*, the 25 mutants were scored for defects in auxin-regulated processes during root and shoot development: root length on medium supplemented with or without IAA, root gravitropic response, and shoot growth including leaf number, rosette size, shoot length, number of axillary and cauline shoots during vegetative and reproductive growth stages (Fig. 4b-c, Fig. S9, Supplementary Table 7). 28% of the mutants showed root phenotypes in processes associated with auxin-regulation, 12% showed auxin-related shoot phenotypes and 52% had both root and shoot phenotypes. For leaf number, rosette size, root elongation and gravitropic response, a majority of the mutants affected showed opposite effects on development from loss of function mutants in loci known to promote auxin signalling (13, 25, 26, 27). This supports a negative regulation of auxin responses by the corresponding TFs. Mutation of single genes in the *ARF*^*ClassA*^ regulatory network can therefore affect auxin-dependent development substantially, demonstrating the functional importance of individual nodes of this regulatory network.

Collectively our results reveal a mechanism through which auxin response is mediated not only through asymmetry in the distribution of the hormone itself, but also through coordinated TF-mediated repression of *ARF*^*ClassA*^ expression. Consistent with this scenario, we often observed that members of the *ARF*^*ClassA*^and their regulatory repressors are expressed in complementary domains in the root (Fig. S10).

Despite a general role of Polycomb-mediated gene repression in tissue-specific expression (28), the absence of H3K27me3 at *ARF*^*ClassA*^loci indicates that their regulation does not rely on this epigenetic mechanism. This may be because such a system would not allow the rapid changes in auxin signalling output required to dampen auxin responses and limit them both spatially and temporally. Instead, we have uncovered a system based on the use of transcriptional repressors to modulate expression of constitutively active loci, which offers a more flexible mode of regulation that can constantly adjust auxin responsiveness. Our network is unlike transcriptional regulation networks previously defined in eukaryotes that involve both transcriptional activators and repressors (29). Instead, our network resembles the early scenario for transcriptional regulation via repressors proposed by Jacob and Monod (30), indicating that there is a biological context for the concept of controlling the expression of key developmental regulators almost entirely via transcriptional repression.

## Methods Summary

### Plant material

T-DNA insertion mutants in transcription factors and the *arf8-1*mutant were obtained from NASC. T-DNA accession numbers are listed in Supplementary Table 5.

### Cloning and generation of ARF transcriptional reporter lines

Multisite Gateway cloning technology was used to generate ARF transcriptional reporter lines harbouring DNA sequences both upstream and downstream from the start codon. The promoter fragments were amplified by PCR with sequences: pARF5 −5418 bp to + 134 bp, pARF6 −3255 bp to +197 bp, pARF7 −2973 bp to + 374 bp, pARF8 −5091 bp to + 42 bp, pARF19 −4906 bp to + 452 bp. For ARF5, 6, 8, and 19 the fragments were inserted into pDONR P4-P1R and recombined with 3x mVenus-N7 pDONR211 (containing triple mVenus coding sequences and N7 nuclear localization signal), OCS terminator pDONR P2R-P3 (containing the stop codon followed by a octopine synthase terminator) and pK7m34GW (the destination vector containing kanamycin resistance gene for *in planta*selection) to produce pARF::mVenus constructs. For ARF7, the fragment was cloned into a pCR8/GW/TOPO and recombined with a nuclear-localized mVenusN7, 35S terminator and pK7m34GW to produce pARF7::mVenus construct. Similarly, the shorter promoter fragments were amplified by PCR based on primers designed at the following locations: pARF5 −5418 bp to −1 bp, pARF6 −3255 bp to −1 bp, pARF7 −2973 bp to −1 bp, pARF8 −5091 bp to −1 bp, pARF19 −4906 bp to −1 bp. The fragments were inserted into pDONR P4-P1R and recombined with 3x mVenus-N7 pDONR211, OCS terminator pDONR P2R-P3 and pK7m34GW destination vector to yield pARF^-^intron::mVenus shorter transcriptional reporter lines. The constructs were transformed into *Agrobacterium tumefaciens*C58pMP90 strain by electroporation and then transformed into Col-0 plants by the floral dip method (Clough and Bent 1998).

### Root microscopy

For root microscopy plants were grown on half-strength Murashige and Skoog (MS) medium supplemented with 1% sucrose and 1% agar. The seedlings were grown in 24h light conditions and imaged at 5 or 6 days in light. Plant cell membranes were visualized by staining with 15 µg/ml propidium iodide solution. Roots were examined using a TCS-SP5 confocal microscope (Leica) with excitation at 514 nm and emission at 526-560 nm for mVenus and 605-745 nm for propidium iodide.

### Shoot microscopy

Plants were grown in 8h light/16h dark conditions for 6 weeks and then transferred to 16h light/8h dark conditions for 2 weeks to induce bolting. The bolted shoots were dissected under a stereomicroscope and planted into an Apex Culture Medium (half-strength MS medium supplemented with 1% sucrose, 0.8% agarose, 1x vitamin solution (myo-Inositol 100 mg/L, nicotinic acid 1 mg/L, pyridoxine hydrochloride 1 mg/L, thiamine hydrochloride 10 mg/L, glycine 2 mg/L)), for overnight incubation at 16h light/8h dark conditions. Before microscopy cell membranes were visualized by staining the shoot apexes with 100 µg/ml propidium iodide solution. The shoot apices were examined using a TCS-SP5 confocal microscope (Leica) with excitation at 514 nm and emission at 526-560 nm for mVenus and 605-745 nm for propidium iodide.

### Y1H assay

The yeast one-hybrid assay was conducted according to Gaudinier *et al.* 2011 (23). The overall ARF promoters screened correspond in length and content to the ones used in the construction of the transcriptional reporter lines: pARF5 −5418 bp to + 134 bp, pARF6 −3255 bp to +197 bp, pARF7 −2973 bp to + 374 bp, pARF8 −5091 bp to + 42 bp, pARF19 −4906 bp to + 452 bp. pARF5, pARF8 and pARF19 promoters were divided in two fragments and each was screened separately.

### Transient expression in *Arabidopsis thaliana* protoplasts

In the transient protoplast assay, a reporter and an effector plasmid were co-transformed (Fig. S6). To produce the reporter plasmid, the promoter fragment of the respective ARF corresponding to the one used in the eY1H assay and the ARF transcriptional reporter lines described above were amplified by PCR and recombined with a pDONR P4-P1R plasmid. For the ARF8 promoter a short part of the 35S promoter (−107 to +1) was inserted at position −115 bp. Separately, a construct containing NLS followed by mVenus coding sequence and an octopine synthase (OCS) terminator was cloned into pDONR 211 plasmid. Thirdly, a construct containing the promoter of RPS5a (promoter of the ribosomal protein S5A) driving TagBFP followed by a NLS signal and a nosT terminator were recombined with the pDONR P2R-P3 plasmid. These three plasmids were recombined with a multisite Gateway to yield the final reporter plasmid pARF-NLS-mVenus-term-pRPS5a-TagBFP-NLS-term. An alternative reporter plasmid contained a shorter ARF promoter fragment which contained sequences upstream and lacked sequences downstream of the start codon (corresponding to the transcriptional reporter lines with shorter promoters described above).

To produce the effector plasmid, the RPS5a was cloned into pDONR P4-P1R plasmid. The cDNA of the respective transcription factor without the stop codon was cloned into pDONR 211 plasmid. The construct contained a self-cleaving 2A peptide (Kim *et al.*2011; Trichas *et al.*2008) followed by mCherry coding sequence, a NLS and a nosT terminator and was cloned into the pDONR P2R-P3 plasmid. Finally, these three plasmids were recombined with a multisite Gateway reaction to yield pRPS5a-cDNA-2A-mCherry-NLS-term. An alternative effector plasmid included an activator VP16 domain from the herpes simplex virus fused to the TF cDNA.

For the protoplast assay Col-0 seedlings were grown in shortday conditions (8h light/16h dark) for 37-45 days. Leaves of similar size from the second or third pair were collected and digested in an enzyme solution (1% cellulose R10, 0.25% macerozyme R10, 0.4M mannitol, 10 mM CaCl2, 20 mM KCl, 0.1% BSA, 20 mM MES at pH 5.7) overnight at room temperature. Protoplasts were collected through a 70 micron mesh, washed twice with an ice-cold W5 solution (154 mM NaCl, 125 mM CaCl2, 5 mM KCl, 5 mM glucose, 2 mM MES at pH 5.7) and incubated on ice for 30 min. The protoplasts were then resuspended in the MMG solution (0.4 M mannitol, 15 mM MgCl2, 4 mM MES at pH 5.7) with a final concentration 150 000 cells/ml. 10 µl of each the effector and the reporter plasmid DNA (concentration 3 mg/µl) were mixed with 200 µl of the protoplasts. Immediately, 220 µl of the PEG solution (40 % PEG 4000, 0.2 M mannitol, 0.1 M CaCl2) was added, incubated for 5 min at RT and then washed twice in W5 solution. The protoplasts were resuspended in 800 µl of the W5 solution and incubated for 24 hours in 16h light/8h dark growth chamber. Before imaging, the protoplasts were resuspended in 400 µl W5 solution and subsequently transformed into an 8-well imaging chamber.

A Zeiss 710 LSM confocal microscope was used for imaging the protoplasts (Fig. S6). The sequential scanning was performed with mVenus (excitation at 514, emission at 520-559), TagBFP (excitation at 405 and emission at 423-491), mCherry (excitation at 561, emission at 598-636) and bright-field channels. Z-stacks of several protoplasts were taken. The data was analysed using ImageJ software. The image with the best focus for each protoplast was selected from the z-stack. The nucleus was selected and the mean fluorescence was measured as illustrated in Fig. S6. For the statistical analysis an unpaired t-test was conducted with p < 0.1 considered as statistically significant. The number of replicates was 40 protoplasts. For each ARF-TF interaction 4 or 5 independent experiments were performed (Supplementary Table 4): 2-3 experiments with the standard effector plasmid and 2 experiments with alternative effector plasmid containing VP16 domain.

### Expression analysis with qRT-PCR

Wild-type and mutant seedlings were grown in 24h light conditions on 1/2 MS plates containing 1% sucrose and 1% agar for 7 days. The whole root and the whole shoot parts of the seedlings were collected separately. For one root sample, roots from 30 seedlings grown on the same plate were pooled together. For one shoot sample, 8 shoots from seedlings grown on the same plate were pooled together. Three independent replicates per genotype were collected. RNA was extracted using Spectrum Plant Total RNA kit (Sigma-Aldrich). The DNA was removed using TURBO DNA-free kit (Invitrogen). The cDNA was produced using SuperScript VILO cDNA Synthesis kit (Thermo Fisher) with 500 ng RNA. The cDNA was diluted 1:100 before use. The qRT-PCR was performed using Applied Biosystems Fast SYBR Green Master Mix. Expression of TUB4 gene was used as standard. The statistical analysis was performed with Mann-Whitney test with p < 0.1 considered as statistically significant.

### Expression analysis of crosses between ARF transcriptional reporter lines and TF mutants

Mutants of the regulatory transcription factors were crossed with pARF7::mVenus transcriptional reporter line described above. The crosses were selected for the presence of a homozygous pARF7::mVenus reporter construct. The F3 generation wild-type and mutant plants were compared. The plants were grown in 16h light / 8h dark conditions. Root microscopy was performed as described above.

### Shoot phenotype analysis of the TF mutants

25 T-DNA insertion mutants and the wild-type Col-0 were grown in 8h light/16h dark conditions on soil for 43 days. Leaf number was counted every 3 days starting from day 24. Rosette diameter was measured at 43 days. After 43 days of growth in the above conditions, the plants were transferred to 16h light/8h dark conditions to induce bolting. The following parameters were measured at 21 and 27 days in the 16h light/8h dark conditions: length of the main stem, number of cauline branches growing from the main stem, number of axillary branches growing from rosette (the main stem not included). The number of replicates per genotype was 12 plants. For the statistical analysis an unpaired t-test was conducted with p ≤ 0.05 considered as statistically significant.

### Root phenotype analysis of the TF mutants

For root length measurement and for gravitropic analysis plants were grown on ½ MS medium supplemented with 1% agar in 12h light/12h dark conditions. For root length analysis, plants were grown either on medium lacking IAA or supplemented with 10 µM IAA. Images were taken at 5 and 15 days in light and the root length was measured. The number of replicates per genotype was at least 26 plants without IAA and 15 plants with IAA. For the gravitropic response, plants were grown for 5 days, then turned at a 90°C angle and the images of the root gravitropic growth were taken every 1 hour for the next 12h hours in the dark with the infrared camera. The number of replicates per genotype was at least 26 plants. Statistical analysis was done with one-way analysis of variance (ANOVA) test with p ≤ 0.05 considered as statistically significant.

### *In silico* analysis

Expression of TFs in the root and the shoot apical meristems was analysed using cell-specific expression profiles from Brady *et al.* 2007, Yadav *et al.* 2009 and Yadav *et al.* 2014 datasets. Overrepresentation of TF gene families was analysed for families represented by two or more members in the network. The number of gene family members in the network was compared to the total number of genes from this family in the TF library. Involvement of TFs in specific developmental processes (development, biotic and abiotic stress) was analysed based on literature description.

### Chromatin state analysis

Binary data on H3K27me3- and H3K4me3 marked genes and chromatin accessibility regions were retrieved from multiple datasets covering a range of tissues and developmental stages. For each dataset, at least two biological replicates were considered and only the presence of a given *ARF* in both gene lists was scored as a positive association with a chromatin mark or an accessible region.

Datasets used for chromatin marking analysis were: H3K27me3 (Roudier et al. 2011; Oh et al. 2008; Willing et al. 2015; Deal et al. 2010; You et al. 2017) and H3K4me3 (Roudier et al. 2011; Oh et al. 2008; Lafos et al. 2011; Willing et al. 2015; You et al. 2017).

Datasets used for chromatin accessibility analysis were: DNase I hypersensitive sites (Gene Expression Omnibus (GEO) database GSM1289358, GSM1289362 and GSM1289374; Sullivan et al. 2014), FANS-ATAC-defined accessible regions (GEO GSM2260231, GSM2260232, GSM2260235, GSM2260236; Lu et al. 2017) and ATAC-defined transposase hypersensitive sites (Maher et al. 2018; Sijacic et al. 2018). For each chromatin accessibility dataset, the presence of at least one accessible region within the *ARF* gene and up to 1 kb upstream of its transcription start site was scored using *ad hoc* scripts. Visualization of epigenomic data was carried out using the IGV software (James T. Robinson, Helga Thorvaldsdóttir, Wendy Winckler, Mitchell Guttman, Eric S. Lander, Gad Getz, Jill P. Mesirov. Integrative Genomics Viewer. Nature Biotechnology 29, 24–26 (2011), Helga Thorvaldsdóttir, James T. Robinson, Jill P. Mesirov. Integrative Genomics Viewer (IGV): high-performance genomics data visualization and exploration. Briefings in Bioinformatics 14, 178-192 (2013).

## Acknowledgements

We thank Jonathan Legrand, Fabrice Besnard and Antoine Larrieu for their help with the data analysis and the statistics; Giuseppe Castiglione for help with root phenotyping and Dolf Weijers for ARF transcriptional reporter lines. This work was supported by ANR-2014-CE11-0018 grant (Serrations) to T.V; a Royal Society University Research Fellowship to A.B.; Aux-ID CNRS PICS grant; a joint INRA/University of Nottingham PhD grant to J.T; an HHMI Faculty Scholar fellowship to S.M.B.

## Author Contribution

J.T., C.S. G.-A., M.B., F.R, S.B., A.B. and T.V. designed experiments; J.T., J.H., C.S. G.-A., S.L., G.B., S.P., A.-M. B., M.E. S. performed experiments; all authors were involved in data analysis; and J.T., A.B. and T.V. wrote the manuscript with inputs from all authors.

